# Melanin Suppresses Aβ Aggregation and Toxicity

**DOI:** 10.64898/2026.09.01.748646

**Authors:** Alexander Röntgen, Jan Lukas Heise, Michele Vendruscolo, Zenon Toprakcioglu

## Abstract

The aggregation of amyloid-β (Aβ) peptides into insoluble deposits is a characteristic hallmark of Alzheimer’s disease (AD) and related neurodegenerative disorders. While AD is the most common cause of dementia, there are currently no disease-modifying treatments which are both affordable and adverse-free. In this study, we report that melanin, a pigment which is commonly found in nature and is abundant in parts of the human brain, suppresses the aggregation of the 42-amino acid Aβ variant (Aβ42). Using biophysical and biochemical techniques, we show that melanin delays Aβ42 aggregation while also reducing the amount of Aβ42 that converts into aggregates. Using thioflavin T assays paired with chemical kinetics, we characterised the melanin-induced inhibition of Aβ42 aggregation *in vitro.* Using MALDI-MS, we elucidate the molecular basis of this effect by showing that melanin prevents Aβ42 dimerisation. We then demonstrate that melanin also reverts the aggregation process by dissolving pre-formed Aβ42 fibrils. Finally, we show that melanin reduces Aβ42 aggregation and rescues Aβ42 toxicity in an SH-SY5Y neuroblastoma cell model. Our study shows that melanin disrupts the aggregation and cytotoxicity of Aβ42, and suggests that compounds derived from human metabolites may offer promising avenues to combat amyloid formation.

## Introduction

Alzheimer’s disease (AD) is a devastating age-related neurodegenerative disorder that affects more than 50 million people worldwide.^1,2^ AD is characterised by the aggregation of amyloid-β (Aβ) peptides into large proteinaceous deposits, known as amyloid plaques, in the brain of afflicted patients.^3–5^ At the molecular level, Aβ peptides of varying lengths are generated through the sequential proteolytic cleavage of the amyloid precursor protein (APP), a widely expressed transmembrane protein which is highly enriched in neurons.^6,7^ In particular, the 42-amino acid Aβ variant (Aβ42) displays an enhanced propensity to aggregate into amyloid fibrils.^8,9^ This aggregation process consists of several microscopic steps.^5,10,11^ Initially, soluble Aβ monomers undergo primary nucleation and form oligomers. Subsequently, through the addition of further monomers, these oligomers grow into mature amyloid fibrils.^5,10,11^ Aβ aggregates are cytotoxic and facilitate neuronal decline, which ultimately leads to the progressive cognitive impairment typically observed in AD.^3–5^ Besides Alzheimer’s disease, Aβ deposits are also found in other neurological disorders, including cerebral amyloid angiopathy (CAA), Parkinson’s disease (PD), and dementia with Lewy bodies (DLB).^12–14^ As the world population is ageing, the number of patients suffering from these diseases is expected to rise, thus further increasing the already high economic and healthcare burden caused by these disorders.^1,15^ At the same time, disease-modifying treatments for AD and related disorders are still not commonly adopted, a situation that at least in part is linked to an incomplete understanding of the biological factors that modulate the Aβ aggregation process. Consequently, understanding the pathogenesis of AD and finding molecules that effectively inhibit Aβ aggregation are subject to ongoing research.^16^

In the past years, extensive efforts have been invested in developing pharmacological interventions that prevent Aβ aggregation or elicit the removal of existing Aβ aggregates.^16–19^ However, the development of amyloid-targeting drugs is complicated because these molecules need to cross the tightly controlled blood-brain barrier while minimising adverse effects in patients. Additionally, the brain is a highly complex environment in which myriads of cellular components can interact with amyloid proteins and the administered drug molecules, which may hamper the desired therapeutic effect. Thus, attention in amyloid research has recently been drawn to metabolites that are endogenous to the human brain. Specifically, studies aim to understand how the molecular properties of human brain metabolites could be harnessed to mitigate amyloid formation and the ensuing neurodegenerative processes.^20–23^

In this study, we explore the action of melanin, a polymeric pigment that is ubiquitous in nature and abundant in parts of the human body. In peripheral human tissues, melanin is produced by specialised cells termed melanocytes and exerts diverse physiological functions, including skin pigmentation, the protection of DNA from ultraviolet radiation, antioxidative effects by scavenging free radicals, and the chelation of heavy metal ions.^24,25^ Furthermore, melanin is produced directly in the human brain and its levels increase with age. Melanin is particularly concentrated in catecholaminergic neurons, where it is thought to protect neuronal viability and firing capacity by acting as a sink for neurotoxic oxidation products of catecholamine neurotransmitters.^26,27^ To date, a number of studies suggest that changes in melanin are linked to the pathogenesis of neurodegenerative disorders. For instance, melanin may become cytotoxic when cellular storage thresholds are exceeded, or when it is released into the extracellular space upon neuronal death, potentially contributing to the degeneration of brain regions early affected in AD and PD.^27^ Moreover, parallels between neuronal melanin and Aβ have been highlighted, including their involvement in both neurotoxic and neuroprotective pathways.^28^ Furthermore, melanin granules have been shown to interact with amyloid deposits termed Lewy bodies in PD patient brains, and the demise of melanin-containing dopaminergic neurons is a well-documented hallmark in PD and DLB.^29,30^

However, despite emerging evidence indicating that melanin is linked to neurodegenerative diseases, the relationship between melanin and amyloid formation has not yet been fully elucidated. To address this gap, we perform a detailed characterisation of the effect of a synthetic construct of melanin on the aggregation of the Aβ42 peptide, using a combination of biophysical, biochemical, and cell-based approaches. At first, we show that melanin delays and reduces the aggregation of Aβ42 using thioflavin T (ThT). Subsequently, we characterise the resulting aggregates using Fourier-transform infrared spectroscopy (FTIR) and transmission electron microscopy (TEM). Using matrix-assisted laser desorption/ionisation mass spectrometry (MALDI-MS), we demonstrate that melanin acts early in the aggregation process by interfering with Aβ42 dimer formation. We then demonstrate that melanin efficiently dissociates pre-formed Aβ42 aggregates. Lastly, we show that melanin inhibits Aβ42 aggregation and rescues Aβ42-induced cytotoxicity in SH-SY5Y neuroblastoma cells.

Taken together, our work demonstrates that melanin is an inhibitor of Aβ42 aggregation, highlighting how compounds derived from human brain metabolites are promising templates for designing modulators of amyloid formation. These insights are particularly relevant given that we currently lack effective treatments to intervene against AD.

## Materials and methods

### Recombinant production and purification of Aβ42

Aβ42 peptides were expressed from a pT7 plasmid in BL21 (DE3) Gold *Escherichia coli* bacteria and were purified based on previously established protocols.^31,32^ The recombinant peptides were eluted in 50 mM Tris-HCl, pH 7.4, using size exclusion chromatography, and the solution was finally aliquoted, snap-frozen in liquid nitrogen, lyophilised, and stored at –80 °C until further use.

### Melanin

Melanin powder (#466430010; ThermoFisher Scientific) was resuspended in 50 mM Tris-HCl, pH 7.4, to a final concentration of 250 µM. The solution was then stirred for 1 min and sonicated in an Ultrawave U100H water bath for 1 min. This procedure was repeated once, and the resulting solution was filtered through a 0.22 µm syringe filter (Millex-GP, Sigma-Aldrich). Because parts of the melanin suspension were filtered out in this last step, the final working concentration of melanin was lower than the concentrations indicated in this study, which are calculated based on the powder that was weighed in.

### Kinetic assays

To assess the aggregation of Aβ42 with melanin, 1 µM Aβ42 was mixed with 50 µM thioflavin T (ThT) and varying concentrations of melanin (1-50 µM) in 50 mM Tris-HCl, pH 7.4. The mixtures were transferred onto Corning 96-well Half-Area Black with Clear Flat Bottom Polystyrene Non-Binding Surface Microplate at 100 µL per well, and the plates were covered using Thermowell Sealing Tape (Corning). ThT fluorescence was then measured over time on a FLUOstar Omega plate reader (BMG Labtech) at 37 °C under quiescent conditions. The aggregation reactions were performed with at least three replicates, and the ThT fluorescence data were normalised to minimum and maximum values. Aggregation half-times were calculated as the timepoints at which the normalised ThT intensity reached 0.5, and median traces were chosen for representation and for the determination of kinetic rate constants using AmyloFit.^33^ To test Aβ42 disaggregation by melanin, amyloid fibrils were generated by incubating 5 µM Aβ42 for one day as described above. After this, the fibrils were recovered from the plate and harvested by centrifugation (1 h, 21,100 *g*, room temperature). Subsequently, the fibrils were sonicated in an Ultrawave U100H water bath for 5 min, diluted to 1 µM and mixed with ThT and different concentrations of melanin, and incubated on the plate reader as described above.

### SDS-PAGE

Samples from the aggregation plate were recovered and centrifuged (1 h, 21100 *g*, room temperature) to separate insoluble from soluble protein species. The supernatant was removed, and the pellet was boiled in 1x NuPAGE LDS sample buffer for 5 min at 95 °C. Subsequently, the samples were run on 4-12% BisTris NuPage using 1x NuPAGE MES SDS Running Buffer and SeeBlue Plus2 Pre-Stained Protein Standard (Thermo Fisher Scientific). Protein bands were visualised using InstantBlue Coomassie Protein Stain (Abcam), and images of the gels were acquired on the ChemiDoc MP Imaging System (BioRad). Finally, the intensities of the protein bands were quantified using Fiji.^34^

### Fourier-transform infrared spectroscopy

Fourier-transform infrared (FTIR) spectroscopy was performed using a Vertex 70 FTIR spectrometer (Bruker) with a *Diamond ATR* unit and a deuterated lanthanum α-alanine-doped triglycine sulphate detector. Aβ42 aggregation reactions were performed as described before using a plate-reader based aggregation assay, except that no ThT was used in these samples. The samples were recovered from the plate, centrifuged, and resuspended in MQ water. 5 μL of the solution were deposited on the prism and FTIR spectra were acquired in the range of 4000-900 cm^-1^.

### Transmission electron microscopy

Samples from aggregation reactions were deposited on glow-discharged 3-mm 300-mesh carbon-film copper grids (EM Resolutions Ltd). First, 3 µL of sample was spotted for 40 s and then blotted using Whatman paper. The sample was then negatively stained with 3 µL 2% (w/v) uranyl acetate for 40 s, blotted again with Whatman paper, and air-dried. Finally, micrographs were acquired on a Talos F200X G2 transmission electron microscope (Thermo Fisher Scientific).

### MALDI-MS

Samples were prepared in 50 mM Tris-HCl, pH 7.4, on ice, containing a final concentration of 5 µM Aβ42 with or without 125 µM melanin. The samples were mixed with matrix solution (75% (v/v) of 20.3 mg/mL 2,5-DHAP in EtOH + 25% (v/v) of 18 mg/mL diammonium hydrogen citrate (DAC) in water) and 2% trifluoroacetic acid (TFA) in a 1:1:1 ratio. 2 µL per mixture were then spotted on polished steel plates and dried under vacuum. Spectra were acquired on an ultrafleXtreme MALDI-TOF/TOF mass spectrometer (Bruker) using linear mode with positive polarity. Finally, spectra were baseline-subtracted, smoothened, and exported using flexAnalysis (Bruker).

### Cell assays

SH-SY5Y neuroblastoma cells were grown in DMEM/F-12+GlutaMAX supplemented with 10% (v/v) fetal bovine serum (FBS; Gibco) on 75 cm^2^ tissue culture flasks (Greiner Bio-One) at 37 °C and 5% CO^2^. For use in experiments, cells were detached from the maintenance plate and seeded at the appropriate density on 96-well plates (Greiner Bio-One). For this, the growth medium was discarded, and the cells were rinsed with 10 mL PBS –CaCl_2_, –MgCl_2_ (Gibco). The cells were then incubated with trypsin-EDTA for 5 min at 37 °C. The cell suspension was recovered from the plate and mixed with growth medium up to a total volume of 10 mL. The cells were harvested by centrifugation (5 min, 300 *g*, room temperature), the supernatant was discarded, and the cell pellet was resuspended in fresh growth medium.

For the assessment of Aβ42 aggregation, 30,000 cells per well were seeded in growth medium on 96-well plates and incubated for 24 h at 37 °C and 5% CO_2_. The cells were then treated with pure RPMI medium (medium control), or RPMI medium containing 1 µM Aβ42 with increasing concentrations of melanin, for 6 h at 37 °C and 5% CO_2_. For cell treatments, Tris buffer, protein and melanin pre-dilutions were sterile-filtered (0.22 µm) under the cell culture hood. After this, wells were stained with Amytracker 680 (1:5000; Ebba Biotech) and 0.5 µM CellTracker Violet BMQC Dye (Thermo Fisher Scientific) for 15 min at room temperature, rinsed with PBS, and imaged on a Leica Stellaris 5 confocal microscope.

For the assessment of cytotoxicity, 10,000 cells per well were seeded in growth medium on 96-well plates and incubated for 24 h. The cells were then treated either with pure growth medium (medium control), or growth medium containing Aβ42 (50-100 nM) with increasing concentrations of melanin, for 24 h at 37 °C and 5% CO_2_. Subsequently, the medium was aspirated and replaced with 10% (v/v) MTT solution (Abcam) in RPMI medium for another 4 h. The medium was again aspirated, and the formazan product was solubilised using cell lysis solution (Abcam) and shaking at 500 rpm and 37 °C for 15 min on a PHMP Grant-Bio Thermoshaker. Finally, the absorbance at 570 nm was measured on a CLARIOstar plate reader (BMG Labtech), the values were baseline-subtracted and normalised to medium control.

## Results

### Melanin delays and reduces Aβ42 aggregation

First, we investigated the effect of melanin (acquired as a synthetic construct from Thermo Fisher Scientific) on the aggregation kinetics of Aβ42. To this end, we mixed 1 µM monomeric Aβ42 with increasing concentrations of melanin and monitored the aggregation reaction by measuring the fluorescence of the amyloid-binding dye thioflavin T (ThT) (**Figure 1A**). ThT binds to β-sheet structures, and the ThT fluorescence intensity is directly correlated with the amount of aggregate mass.^35,36^ The control experiment, which contained Aβ42 but no melanin, displayed a typical sigmoidal curve with an aggregation half-time of ∼1.5 h. In conditions which contained increasing melanin concentrations alongside Aβ42, the aggregation of Aβ42 was progressively delayed and the ThT fluorescence intensity was gradually reduced. Moreover, at melanin:Aβ ratios greater than 15:1, the fluorescence traces barely increased above their baseline values, which suggested a complete inhibition of Aβ42. To further explore this effect, we plotted the ThT fluorescence values in the final plateau phase of each aggregation reaction (**Figure 1B**). Even at the lowest melanin:Aβ ratio tested (5:1), the ThT fluorescence intensity at the reaction endpoint was reduced compared to the Aβ42 control reaction.

**Figure 1.**
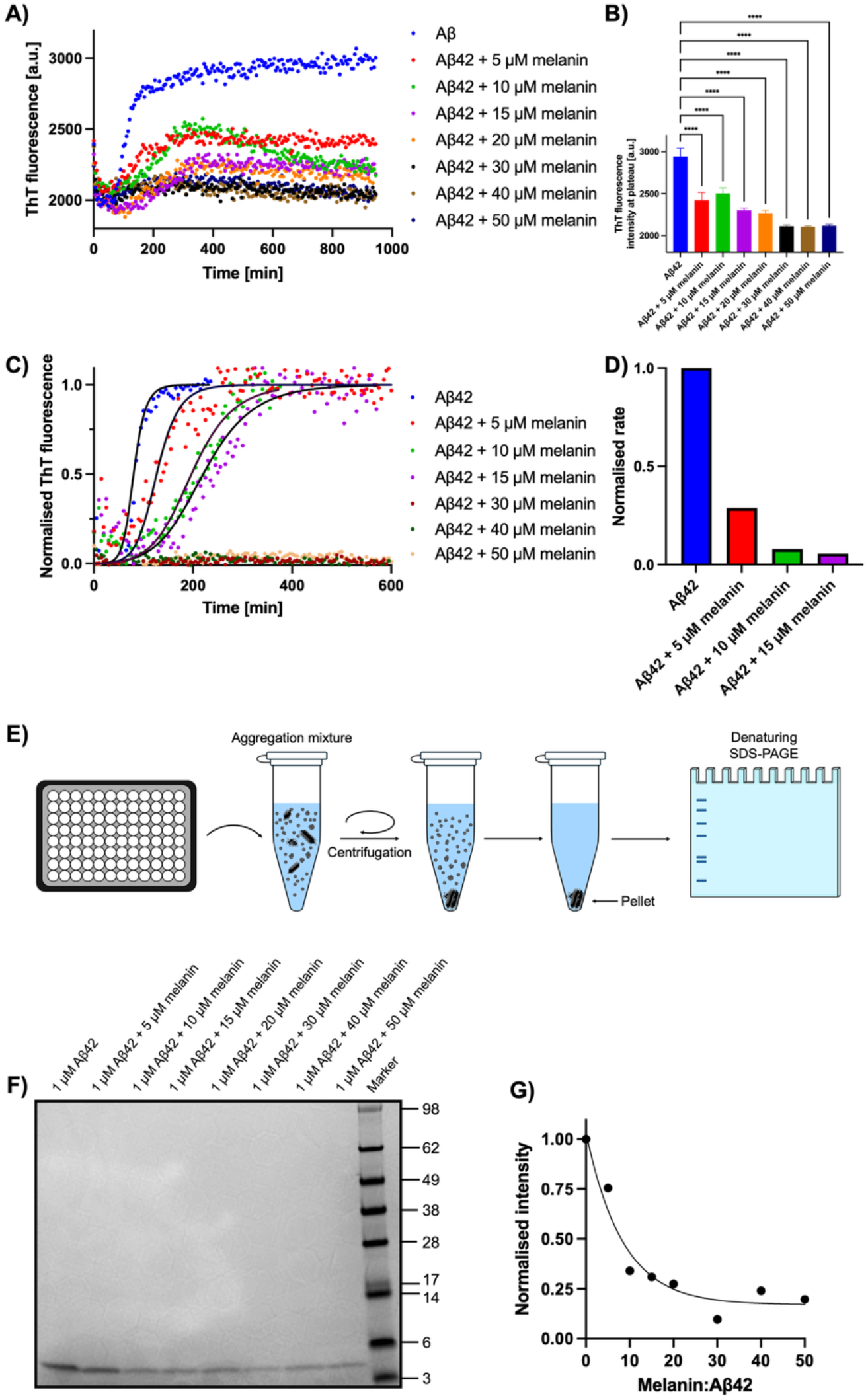
Melanin inhibits Aβ42 aggregation *in vitro*. **(A)** Aggregation traces of Aβ42 with increasing concentrations of melanin. **(B)** Raw ThT values during the plateau phase of the aggregation reaction for the different systems tested in (A). **(C)** Normalised aggregation traces including kinetic fits (solid lines) from raw data in (A), which were performed using AmyloFit.^33^ **(D)** Combined kinetic rates of aggregation for different Aβ42:melanin systems normalised to Aβ42 control. **(E)** Schematic overview of the experimental workflow using centrifugation and SDS-PAGE. The aggregation mixtures were recovered from the aggregation plate and centrifuged to separate insoluble, aggregated species (pellet) from soluble species (supernatant). The supernatant was removed, after which the content of the pellets was denatured, separated by size using SDS-PAGE, and visualised by Coomassie stain. The schematic overview is adapted from^37^. **(F)** Gel image of the aggregation reaction of Aβ42 with increasing amounts of melanin. **(G)** Quantification of the protein band intensities normalised to the Aβ42 control.

To understand the effect of melanin on the aggregation kinetics of Aβ42 in greater detail, we normalised the aggregation traces from **Figure 1A** and fitted them using AmyloFit (**Figure 1C**).^33^ We then used the kinetic fits to calculate the normalised rate of the aggregation reaction (**Figure 1D**). Compared to the Aβ42 control reaction, the combined kinetic rate was decreased by 72% at 5:1 (melanin:Aβ), 92% at 10:1 (melanin:Aβ), and 95% at 15:1 (melanin:Aβ). These results show the degree to which melanin inhibits Aβ42 aggregation (**Figure 1D**).

Taken together, these data indicate that melanin exerts a dual effect on Aβ42. Melanin interacts with Aβ42 in a way that delays aggregation, and it also reduces the amount of the peptide that converts into amyloid aggregates.

To verify that melanin does not merely interfere with the ThT fluorescence signal, we independently determined the fraction of aggregated Aβ42 using SDS-PAGE (**Figure 1E**). First, we recovered the protein from the aggregation reactions and centrifuged the samples to separate the insoluble Aβ42 aggregates from soluble monomeric or oligomeric species. We then denatured the protein pellets containing the aggregated Aβ42 protein, subjected the samples to SDS-PAGE, and stained the protein bands with Coomassie (**Figure 1F**). Using this protocol, we confirmed that the fraction of Aβ42 that is finally aggregated and thus partitions into the insoluble pellet after centrifugation is reduced in the presence of melanin. We quantified this effect by measuring the intensities of the peptide bands in the gel image (**Figure 1G**). This analysis indicates that, above a melanin:Aβ ratio of 5:1, the band intensities are ∼25% compared to the Aβ42 control reaction. Thus, the results from this experiment are consistent with the reduction in aggregation observed using ThT (**Figure 1A-D**).

Overall, our results indicate that melanin inhibits Aβ42 aggregation in a two-fold manner. At intermediate melanin:Aβ ratios (up to 15:1), there is a marked delay in the rate of Aβ42 aggregation and gradual reduction in the total fibrillar amount. At higher melanin:Aβ ratios, Aβ42 aggregation is suppressed to an extent that sigmoidal aggregation traces are no longer observed by measuring ThT fluorescence.

We next considered whether the interaction of melanin with Aβ42 also affects the structure of the resulting Aβ42 aggregates. For this, we used a similar strategy as before, by incubating Aβ42 with different concentrations of melanin and recovering the end product of the aggregation reactions by centrifugation. We then analysed these samples using Fourier-transform infrared (FTIR) spectroscopy to assess the secondary structure of the recovered aggregates (**Figure 2**).

**Figure 2.**
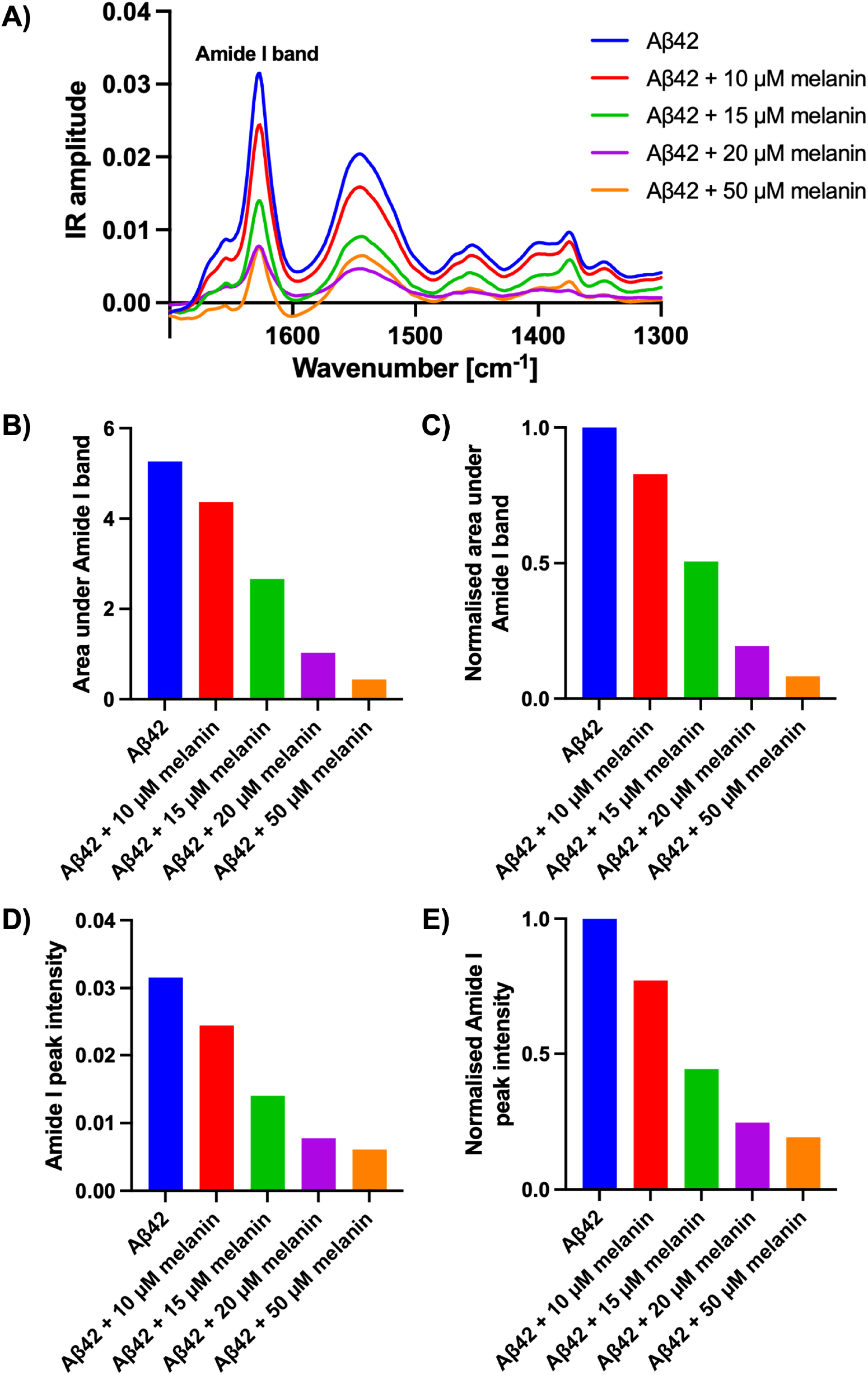
Fourier-transform infrared spectroscopy of Aβ42 aggregated *in vitro* in the presence of melanin. **(A)** FTIR spectra of fibrils resulting from the aggregation of 5 µM Aβ42 formed with increasing ratios of melanin. **(B,C)** Quantification of the area under the Amide I band in absolute values (B) and normalised to the Aβ42 control (C). **(D,E)** Quantification of the Amide I peak intensity in absolute values (D) and normalised to the Aβ42 control (E).

For all conditions, we found that the Amide I band was located around 1625 cm^-1^, which is characteristic for the cross-β architecture of amyloid fibrils (**Figure 2**).^38^ Moreover, the position of the Amide I band remains largely unchanged when increasing amounts of melanin were added to the aggregation reaction. Hence, these data indicate that the aggregates generated through the co-incubation of Aβ42 with melanin exhibited a similar secondary structure content as the fibrils from the Aβ42 control reaction.

We also note that the area under the Amide I peak as well as the peak intensity were reduced with increasing concentrations of melanin. Both of these features are correlated to the amount of β-sheet content in the system, as a larger absolute peak area and intensity corresponds to more sample deposited onto the prism detector of the FTIR spectrometer. This observation suggests that, while the Aβ42 protein that was recovered had a similar secondary structure profile across all conditions tested, the absolute amount of Aβ42 that had converted from monomers into amyloid aggregates was decreased when more melanin was added to the aggregation reaction. Thus, these FTIR data support our previous findings using ThT assays and SDS-PAGE (**Figure 1**), showing that melanin inhibits Aβ42 aggregation and diminishes the fraction of Aβ42 that converts into insoluble, pelletable aggregates.

### Effect of melanin on the morphology of Aβ42 aggregates

We further investigated whether the addition of melanin affected the overall morphology of the resulting Aβ42 aggregates. For this, we spotted samples of the Aβ42 aggregation reactions and imaged them using TEM (**Figure 3**). In this case, we did not separate soluble from insoluble species via centrifugation in order to visualise the different molecular species formed in the aggregation process. As expected, for the Aβ42 control, we observed a dense network of amyloid fibrils, which were a few nanometres in width (**Figure 3A**). In contrast, for the systems containing melanin, a reduced number of fibrils was observed on the electron microscopy grids (**Figure 3B-D**). This reduction in the Aβ42 fibril number was dose-dependent on the melanin concentration present in the aggregation reaction. This observation further corroborates our previous results that melanin interferes with the self-assembly of Aβ42 into amyloid aggregates and reduces the overall fibrillar content.

**Figure 3.**
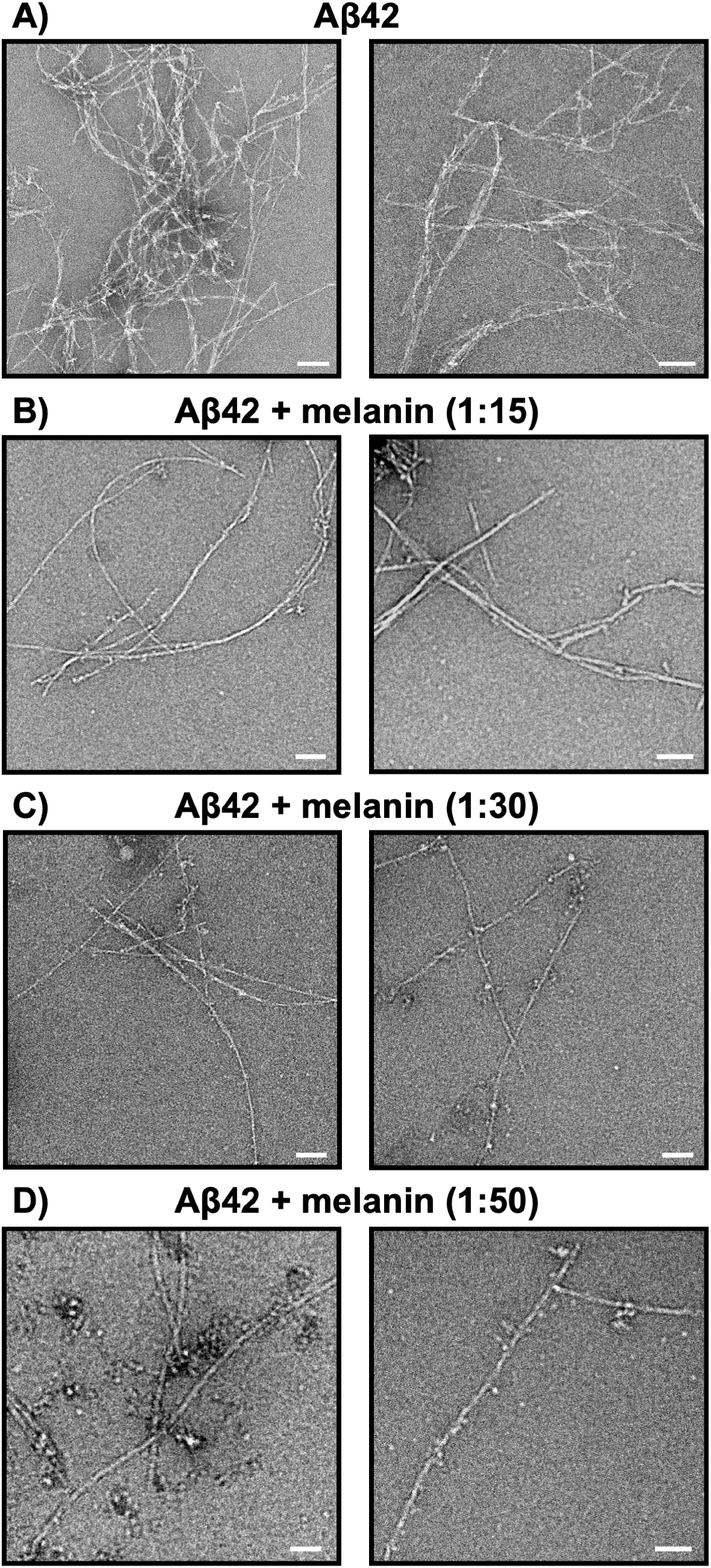
TEM images of Aβ42 aggregates with increasing amounts of melanin. **(A)** Aβ42 alone shows a dense fibrillar network. **(B-D)** Increasing amount of melanin reduces the overall Aβ42 fibrillar aggregate content. Scale bars = 100 nm.

### Melanin suppresses Aβ42 dimer formation

To elucidate the effect of melanin on Aβ42 aggregation in further detail and obtain molecular information on the mechanism of interaction, we next sought to determine which Aβ42 species in the aggregation pathway are predominantly affected by melanin. For this, we explored the possibility that melanin interacts with soluble Aβ42 species and prevents them from converting into mature aggregates. To this end, we incubated monomeric Aβ42 with melanin and employed MALDI-MS to detect low-weight Aβ42 assemblies (**Figure 4A**). In the Aβ42 control experiment, we observed signals for both the Aβ42 monomer and dimer (**Figure 4B**; a magnified view of the Aβ42 dimer peak is shown in the right panel). The dimeric species of Aβ42 reached an intensity of ∼14.5% relative to the monomer peak, which illustrates the propensity of Aβ42 to form oligomers immediately after resuspension.

**Figure 4.**
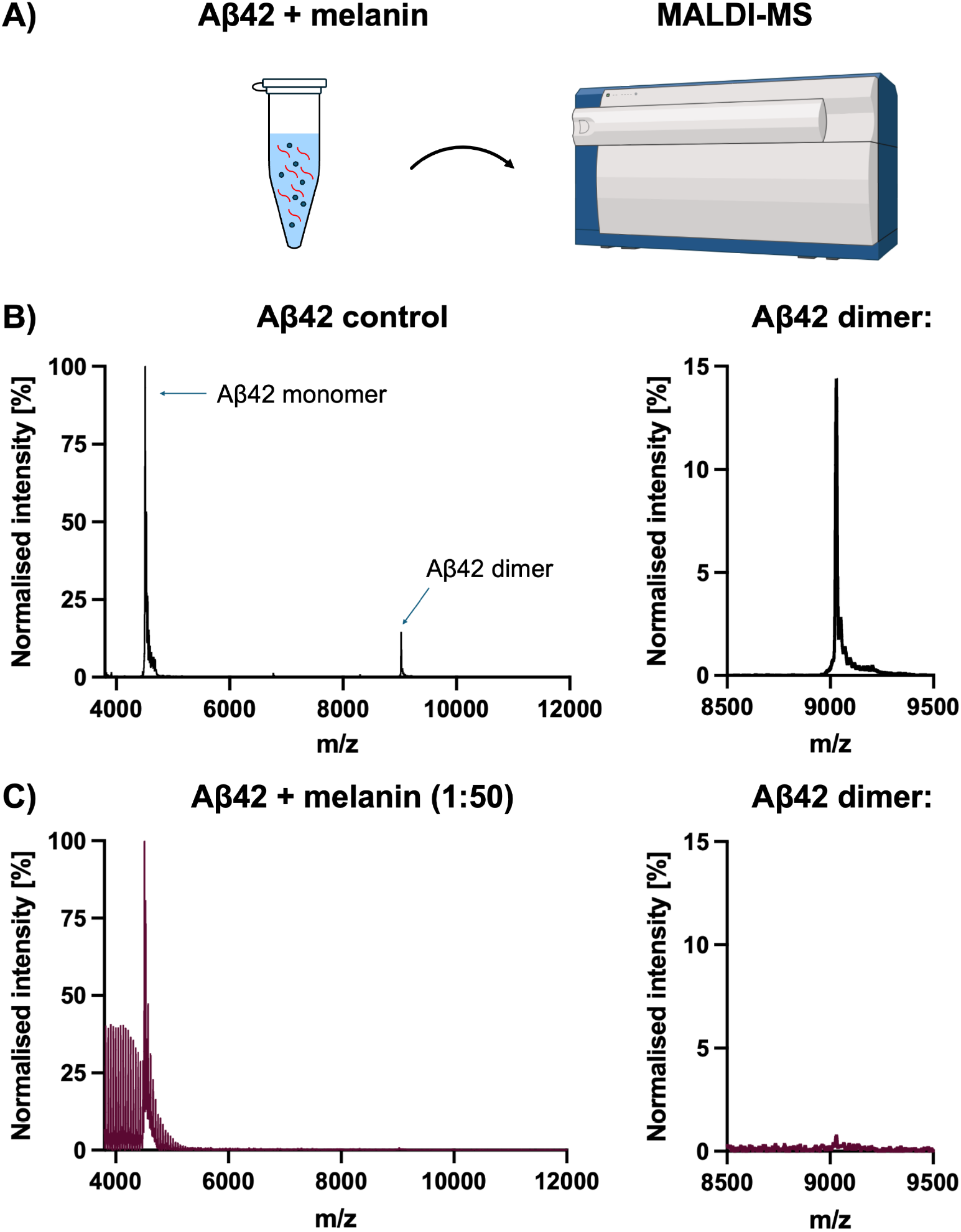
MALDI-MS spectra showing that melanin suppresses Aβ42 dimerisation. **(A)** Schematic overview of the experiment. Elements of this figure have been adapted from biorender.com. **(B,C)** MALDI-MS spectra of 5 µM Aβ42 alone (B) or in mixture with a 50-fold excess of melanin (C). Full spectra are displayed on the left panels, and magnifications showing the Aβ42 dimer are displayed on the right panels.

We next mixed freshly resuspended Aβ42 with melanin and analysed the sample using MALDI-MS (**Figure 4C**). In this sample, a range of additional peaks occurred in the range up to *m/z* 5000. This observation can be linked to the presence of polyethylene glycol in the synthetic melanin preparation. It should be noted that trace amounts of PEG are sufficient to dominate the spectrum due to the high ionisability of PEG.^39,40^ Nevertheless, the Aβ42 monomer peak was well detected (**Figure 4C**). In this spectrum, we found that the Aβ42 dimer was barely present, and merely accounted for ∼0.75% of the signal intensity compared to the monomer peak in the same spectrum, which represents a ∼20 times reduction in the relative signal intensity compared to the Aβ42 control condition. Thus, using MALDI-MS, we confirmed that melanin suppresses the nucleation and formation of Aβ42 aggregates at its earliest stage, *i.e.* the formation of Aβ42 dimers. This observation may be due to Aβ42 monomers interacting with melanin, which renders them unable to form oligomeric nuclei that would eventually elongate into mature amyloid fibrils.

### Melanin promotes the disassembly of pre-formed Aβ42 aggregates

We next considered whether melanin can also affect pre-formed Aβ42 aggregates. For this, we formed Aβ42 fibrils using a plate reader, recovered the aggregates, and mixed them with varying concentrations of melanin. We then added ThT and transferred these mixtures back onto the plate reader to monitor ThT fluorescence over time. In all conditions that contained Aβ42 aggregates and melanin, a decline in ThT fluorescence was observed following the start of the measurement (**Figure 5A**). Subsequently, the ThT fluorescence values approached a stable plateau value over time (**Figure 5A**). In contrast, the control reaction, which contained only Aβ42 and no melanin, did not exhibit an initial decrease and remained stable in its ThT fluorescence levels (**Figure 5A; black curve**).

**Figure 5.**
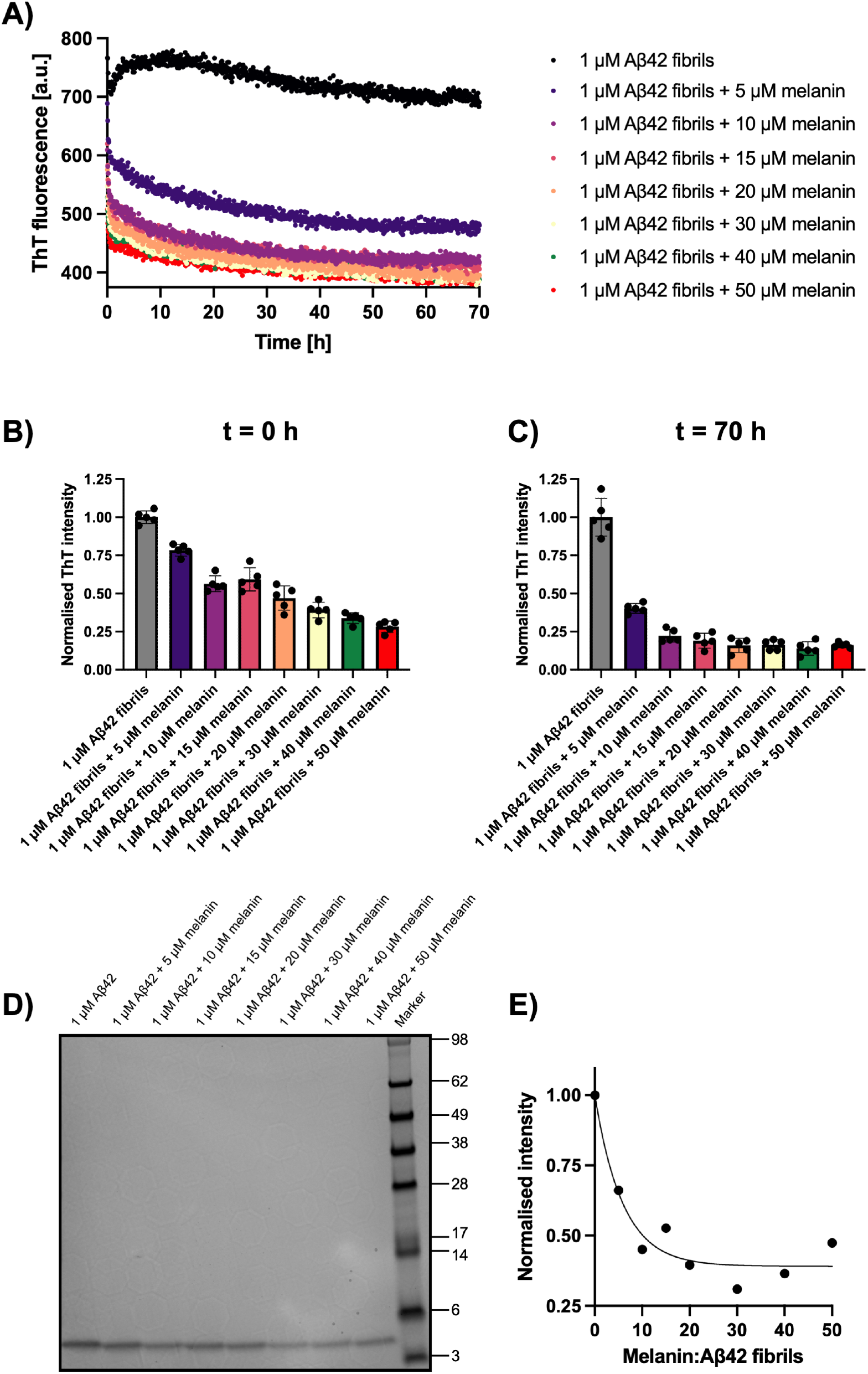
Melanin promotes the disassembly of Aβ42 aggregates. **(A)** ThT traces of Aβ42 fibrils with increasing concentrations of melanin. **(B)** Normalised ThT values at the beginning of the disaggregation reaction (t = 0 h). **(C)** Normalised ThT values at the end of the disaggregation reaction (t = 70 h). **(D)** SDS-PAGE of the pelleted protein recovered at the end of the disaggregation reaction, followed by Coomassie stain. **(E)** Quantification of the protein band intensities normalised to the Aβ42 control.

To further quantify this effect, we normalised the ThT values at the beginning (*i.e.* at t = 0 h) and at the end (*i.e.* at t = 70 h) of the kinetic experiment. For this, we subtracted ThT baseline values, *i.e.* control wells containing only ThT and buffer, from each trace and then divided the values for each condition by the Aβ42 control reaction (**Figure 5B,C**). This normalisation revealed that, at the beginning of the measurement, the ThT fluorescence in all melanin-containing conditions was significantly reduced, suggesting a rapid onset of the melanin-induced disassembly of Aβ42 aggregates (**Figure 5B**). These values were furthermore reduced at 70 h, which reflects a steady decline in fibrillar mass over time (**Figure 5C**).

Additionally, we performed experiments similar to those conducted for **Figure 1**. We centrifuged the end product of the disaggregation reaction, denatured the pellets, subjected them to SDS-PAGE, and stained the resulting protein bands using Coomassie (**Figure 5D**). In line with our ThT experiments, quantification of the protein band intensities revealed a trend whereby the amount of aggregated protein that was recovered in the insoluble fraction after centrifugation was decreased in all conditions containing melanin (**Figure 5E**). Hence, these data demonstrate that melanin does not only prevent Aβ42 aggregation, but it is also capable of eliciting the disassembly of already existing Aβ42 aggregates.

### Melanin reduces Aβ42 aggregation and toxicity on neuroblastoma cells

To explore the effect of melanin on Aβ42 aggregation in a complex molecular environment, we co-incubated SH-SY5Y neuroblastoma cells with Aβ42 and melanin (**Figures 6 and S1**). We then stained the resulting aggregates using Amytracker 680, which increases its fluorescence signal when bound to amyloid structures. Furthermore, considering that Aβ is prone to interacting with lipid bilayers, we added CellTracker to visualise the location of the aggregates relative to the SH-SY5Y cells (**Figures 6A, S1**).^21,41^ Using this approach, we found that monomeric 1 µM Aβ42 self-assembled and formed Amytracker-positive aggregates that colocalised with the neuroblastoma cells (**Figure 6A, red aggregates clearly shown using white arrows**). Moreover, when melanin was added to the cells, far fewer aggregates were formed, and barely any aggregates were observed at the highest melanin concentration (50 µM) (**Figure 6B,C**). This assay extends our previous experiments by showing that melanin exerts its inhibitory effect on Aβ42 not only *in vitro* but also under cell culture conditions.

**Figure 6.**
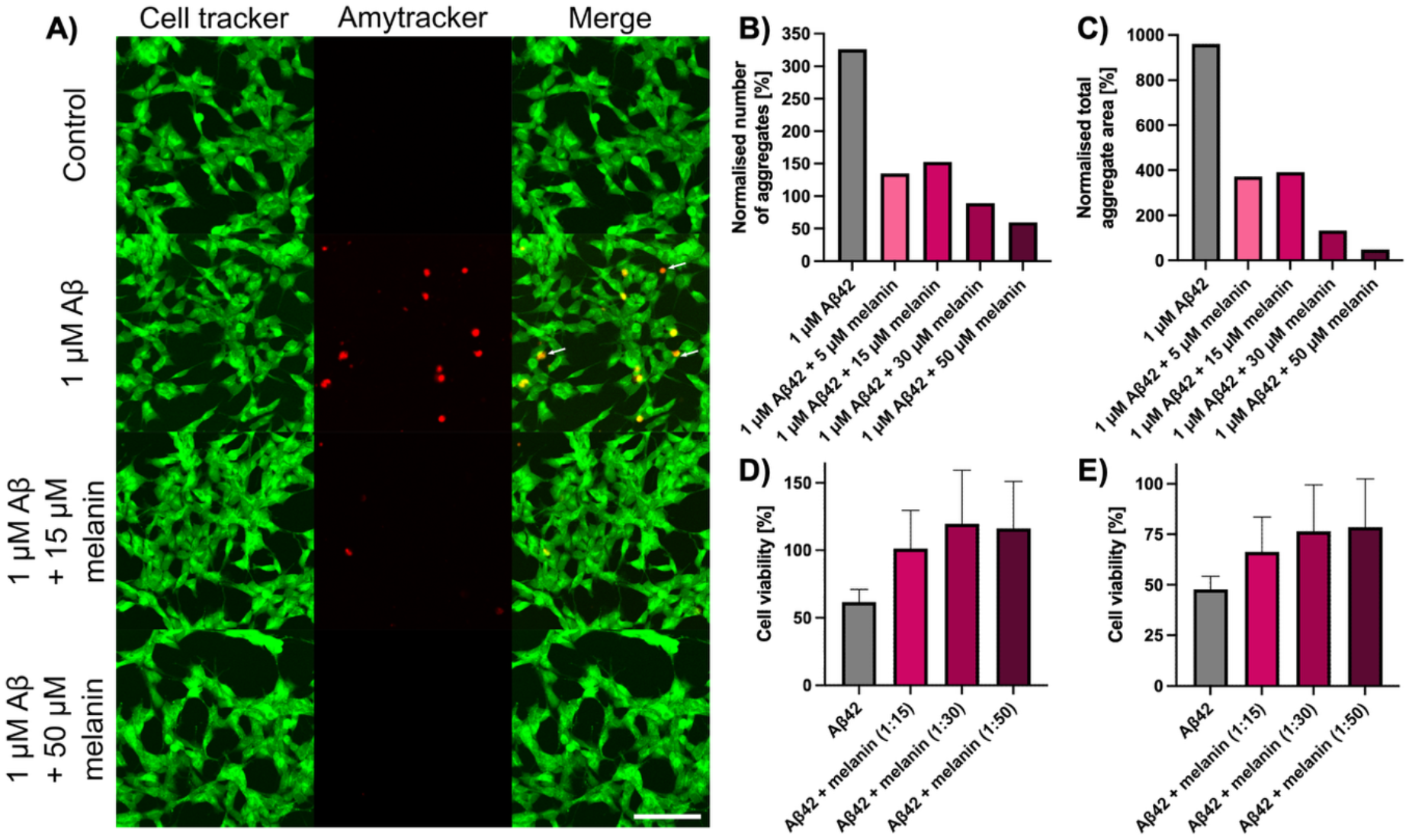
Melanin inhibits Aβ42 aggregation and toxicity in SH-SY5Y neuroblastoma cell culture. **(A)** Representative confocal microscopy images following the incubation of SH-SY5Y cells with Aβ42 and melanin. Aβ42 aggregates were stained with Amytracker 680 (red), and SH-SY5Y neuroblastoma cells were stained with CellTracker (green). Scale bar = 100 µm. **(B,C)** Number and total area of aggregates normalised to medium control. **(D,E)** Cell viability of SH-SY5Y cells following incubation with a fixed concentration of Aβ42 (50 nM (left) or 100 nM (right)) and melanin at varying ratios was measured using MTT. Data are normalised to medium control and are shown as mean ± SEM of independent experiments.

As Aβ42 aggregates are well known to induce cytotoxicity on neurons, we tested whether the melanin-induced reduction in Aβ42 aggregates was correlated with reduced levels of cytotoxicity on neuroblastoma cells. For this, we used an MTT assay to determine the levels of cell viability following incubation with Aβ42 and melanin. We found that the presence of melanin led to an improvement in cell viability. At 50 nM Aβ42, cell viability increased from 62% (Aβ42 control) to 120% (30:1, melanin:Aβ42), while at 100 nM Aβ42, cell viability increased from 48% (Aβ42 control) to 79% (50:1, melanin:Aβ42), representing relative increases by a factor of ∼2.0, and ∼1.7, respectively. As a control experiment, we studied the effect that melanin alone has on the viability of cells. As expected, we found that melanin by itself is completely biocompatible, but notably does not increase cellular viability (**Figure S2**). The increase in cell viability we observed for samples containing Aβ42 and melanin must therefore be due to the fact that melanin reduced the fibrillar content, either by inhibiting Aβ42 aggregation, disassembling Aβ42 fibrils, or through a combination of the two processes.

Thus, our data show that melanin is an inhibitor of Aβ42 both *in vitro* and in cellular systems, and that melanin rescues Aβ42-induced cytotoxicity. Overall, our data are consistent with a model whereby melanin sequesters monomers and potentially other soluble forms of Aβ42, thereby preventing Aβ42 from undergoing aggregation and exerting cytotoxic effects on cells. Taken together, these findings link melanin with the aggregation and toxicity of Aβ42.

## Discussion

We have demonstrated that melanin delays and reduces the aggregation of the Alzheimer’s-associated Aβ42 peptide. The inhibitory effect which we observed prevented most Aβ42 from converting into fibrillar aggregates (**Figures 1 and 2**). Furthermore, structural analysis of the aggregation mixture at the reaction end-point supported a reduction in the formation of mature amyloid fibrils when adding melanin, in a dose-dependent manner (**Figure 3**). Moreover, we showed via MALDI-MS that melanin interacts with Aβ42 at the monomeric level, and consequently acts at the early stages of aggregation by inhibiting Aβ42 dimerisation (**Figure 4**). Melanin not only inhibited Aβ42 aggregation, but it also disassembled pre-formed Aβ42 aggregates (**Figure 5**). Finally, we explored the effect of melanin on cell models and showed that melanin disrupts Aβ42 aggregation and rescues Aβ42 toxicity on neuroblastoma cells (**Figure 6**).

Our biophysical and biochemical results using ThT fluorescence, kinetic analysis, SDS-PAGE, FTIR, TEM and MALDI-MS (**Figures 1-4**) support a model according to which melanin sequesters soluble, monomeric species of Aβ42, which are thus no longer available to participate in the aggregation process. This in turn results in a delay of amyloid aggregation. Overall, this mode of action explains the dual effect we observe, as Aβ42 aggregation is delayed while also the fraction of Aβ42 that eventually converts into aggregates is reduced in the presence of melanin (**Figures 1 and 2**). The range in which we observe these results is where both species are present in similar amounts in terms of mass per volume. Given that the molecular weight of Aβ42 is ∼14 times the molecular weight of the melanin construct used, these species are present in equal mass concentrations at a molar ratio of ∼14:1 (melanin:Aβ). Based on this, a possible explanation for the inhibitory effect of melanin could be that melanin effectively coats monomers or other soluble species of Aβ42, thereby preventing them from interacting with each other and suppressing their aggregation. In addition, melanin itself is known to exist as a polymer.^25^ Hence, melanin polymers could provide surfaces that act as sequestration sites to which Aβ42 monomers or small oligomers may adsorb. With respect to the disassembly of pre-formed Aβ42 fibrils, the above explanation would imply that melanin sequesters monomers or small oligomeric species that dissociate and fragment from fibrils. With progressive spontaneous dissociation of monomers or small soluble species, this process would gradually lead to the dissolution of Aβ42 fibrils. Alternatively, the preferential interaction with melanin could promote the detachment of Aβ42 molecules from fibrils. Of note, the anti-aggregative effects of melanin approaches a maximum at molar ratios between 10:1-15:1 (melanin:Aβ) (**Figures 1 and 5**). The above considerations would explain why molar ratios greater than ∼15:1 (melanin:Aβ) do not lead to a stronger inhibitory or disaggregating effect, as the interaction between the two species becomes saturated. In any case, further investigations and studies will need to be conducted to help determine which moiety of the melanin molecule mediates the inhibitory effect, and to confirm the exact molecular mechanism through which melanin disassembles Aβ42 aggregates.

In the search for compounds that inhibit the aggregation of Aβ, an important advantage of screening metabolites that are endogenous to the human body is that they are already biocompatible. In contrast, other non-human metabolites or organic compounds often need to be subjected to lengthy optimisation procedures to reduce their toxicity. In line with this, previous studies have investigated how the aggregation of amyloid proteins is affected by L-DOPA and dopamine, *i.e.* precursors of melanin, as well as catecholamine-derived metabolites and particles. These results support our findings by showing that catecholamines, too, inhibit amyloid aggregation and lead to the disassembly of amyloid fibrils.^20,22,23^ Even if these compounds will not be used directly as therapeutic agents, they may inform future drug development efforts to design molecules with favourable pharmacokinetic properties for administration to patients while achieving comparable or further improved inhibitory effects on amyloid aggregation.

Among lipid metabolites, retinoic acid (vitamin A), was shown to cause a ∼50% increase in the aggregation half-time at a 10-fold molar excess with Aβ42.^42^ More generally, lipids are an abundant class of biomolecules in the brain and play crucial roles in maintaining the structural and functional integrity of cellular systems. Therefore, lipids have been investigated with respect to modulating the aggregation of Aβ and other amyloidogenic proteins^43^. Gangliosides were found to significantly delay Aβ42 aggregation and even lead to an abrogation of aggregation of the less aggregation-prone Aβ40 variant.^21^ The non-human aminosterol claramine, which has the ability to pass through the blood brain barrier,^44^ has been extensively characterised regarding its effect on Aβ aggregation. In the condensation pathway, claramine was shown to facilitate the transition of Aβ40 into the liquid phase. At low ratios, claramine accelerated Aβ40 aggregation within liquid condensates, while at higher ratios, this effect was reverted and claramine inhibited Aβ40 liquid-to-solid transitions.^45^ Similarly, in the deposition pathway, using plate-reader based aggregation assays, claramine was found to accelerate Aβ42 aggregation kinetics at low ratios but to inhibit Aβ42 aggregation at higher ratios.^46^

Overall, melanin appears to be a human metabolite that reduces the aggregation and cytotoxicity of Aβ. Indeed, melanin has been previously linked to amyloid deposits in the brain.^29,47^ Early structural studies revealed that melanin granules are associated with Lewy bodies, *i.e.* the hallmark deposits in PD brains that consist of amyloid aggregates of the neuronal protein α-synuclein.^29^ Melanin granules and Lewy bodies do not always exhibit clear boundaries, as melanin polymers interact with α-synuclein fibrils.^29^ Conversely, melanin biosynthesis is catalysed by amyloid fibrils formed of the protein Pmel17.^47^ Taken together, melanin and amyloid formation are linked at the molecular level and exert reciprocal modulation. This association supports the view that melanin has an affinity to interact with amyloidogenic proteins and emphasises that melanin could play an important role in modulating amyloid formation in the brain.

We should note that melanin exists in various forms in the human body, including neuromelanin in the brain, as well as eumelanin and pheomelanin in peripheral tissues such as human hair and skin. These variant forms of melanin undergo different biosynthetic pathways, and the resulting polymers exhibit distinct molecular structures.^25^ In this study, we started by using a synthetic construct which was readily available from a commercial source. Consequently, it will be key to understand the effects of naturally occurring forms of melanin. Future studies should address the influence of neuromelanin on amyloid deposition in the brain, while other forms of melanin should be explored for their potential interaction with peripherally depositing amyloid proteins.

Similarly, it has recently become clear that most proteins, including those that form amyloid aggregates, exist as an array of interrelated protein variants instead of single, well-defined molecular species.^48,49^ These protein variants, called proteoforms, exhibit vastly different biophysical properties, including their propensities to undergo amyloid aggregation. This phenomenon has been reported for Aβ and α-synuclein.^9,50,51^ Furthermore, it is now well-established that most amyloid-forming proteins, including Aβ and α-synuclein, do not act in isolation, but rather form co-pathologies in the brains of afflicted patients.^12,13^ Consequently, many neurodegenerative disorders form a disease spectrum instead of representing distinct pathological entities. In particular, Aβ peptides have been shown to promote the nucleation of α-synuclein condensation and aggregation.^37,52,53^ In the future, it will be valuable to disentangle the effects of melanin interacting with complex mixtures of amyloid proteins and their alternative proteoforms. Overall, future studies should investigate the possible effects of melanin in the pathogenesis of neurodegenerative diseases in order to obtain a fundamental understanding of the underpinning molecular mechanisms that govern these misfolding disorders.

## Acknowledgements

A.R. acknowledges funding from the TWIN2PIPSA programme under the grant agreement number 101079147. Z.T. acknowledges funding from the Ron Thomson Research Fellowship in Alzheimer’s Disease, Pembroke College, Cambridge. The authors acknowledge the EPSRC Underpinning Multi-User Equipment Call (EP/P030467/1) for funding of the electron microscopy facility (Yusuf Hamied Department of Chemistry, University of Cambridge).

## Supplementary Information

**Figure S1.**
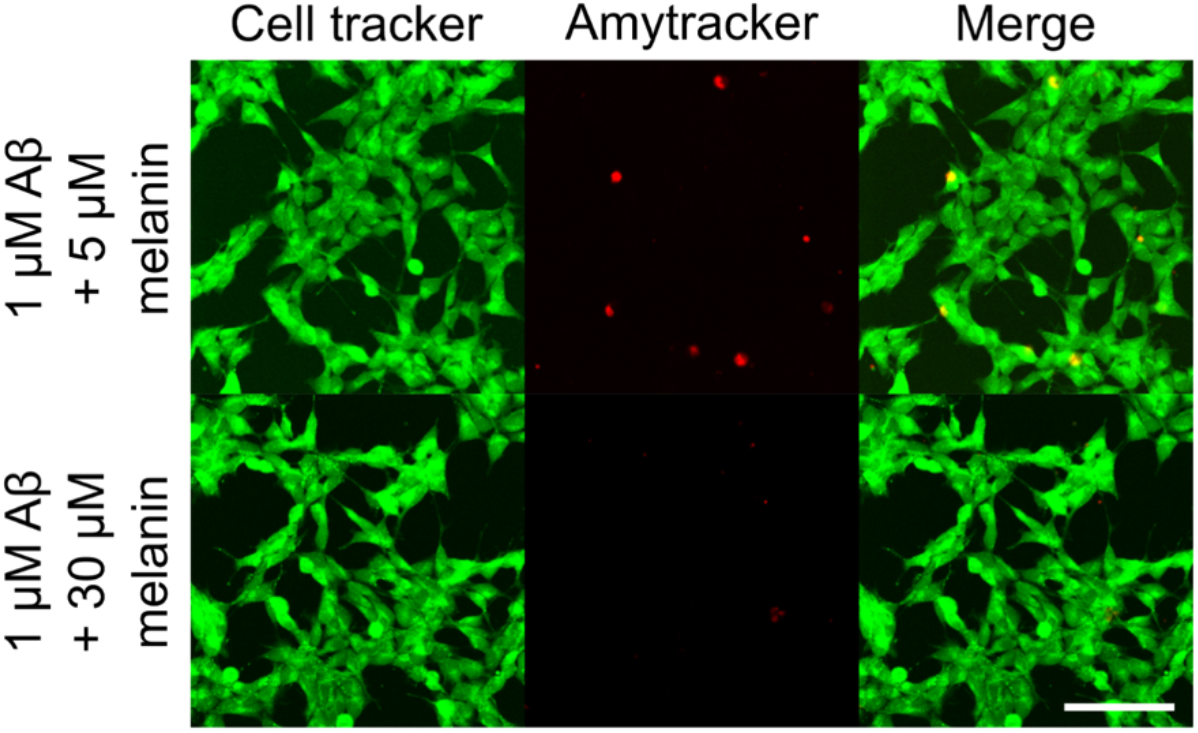
Additional conditions tested in cellular assay. Scale bar = 100 µm.

**Figure S2.**
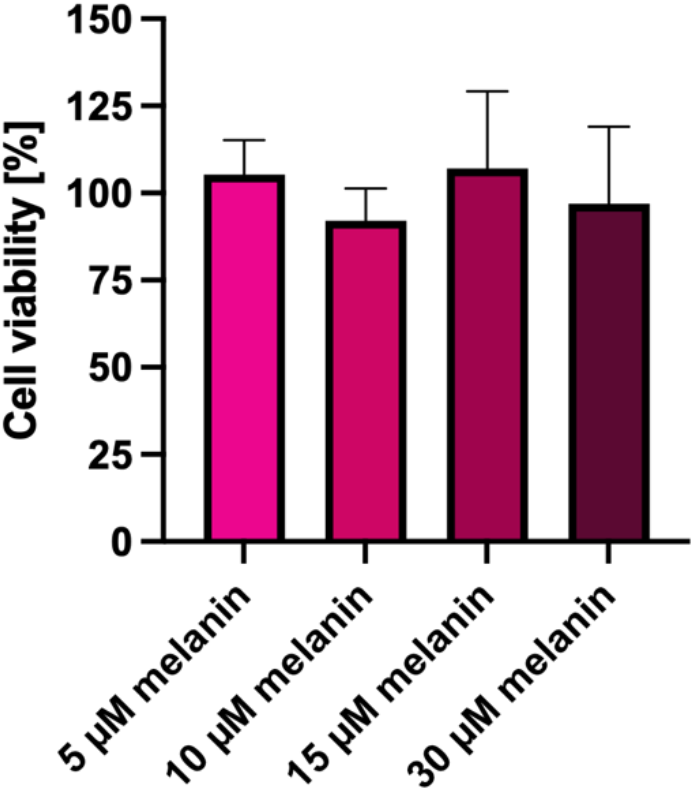
**Melanin alone does not change SH-SY5Y cell viability.**

